# Diffusion barriers imposed by tissue topology shape morphogen gradients

**DOI:** 10.1101/2024.05.01.592050

**Authors:** Gavin Schlissel, Miram Meziane, Domenic Narducci, Anders S. Hansen, Pulin Li

## Abstract

Animals use a small number of morphogens to pattern tissues, but it is unclear how evolution modulates morphogen signaling range to match tissues of varying sizes. Here, we used single molecule imaging in reconstituted morphogen gradients and in tissue explants to determine that Hedgehog diffused extra-cellularly as a monomer, and rapidly transitioned between membrane-confined and -unconfined states. Unexpectedly, the vertebrate-specific protein SCUBE1 expanded Hedgehog gradients by accelerating the transition rates between states without affecting the relative abundance of molecules in each state. This observation could not be explained under existing models of morphogen diffusion. Instead, we developed a topology-limited diffusion model in which cell-cell gaps create diffusion barriers, and morphogens can only overcome the barrier by passing through a membrane-unconfined state. Under this model, SCUBE1 promotes Hedgehog secretion and diffusion by allowing it to transiently overcome diffusion barriers. This multiscale understanding of morphogen gradient formation unified prior models and discovered novel knobs that nature can use to tune morphogen gradient sizes across tissues and organisms.

## Introduction

Morphogens are secreted signaling molecules that form concentration gradients to establish positional information during embryonic development. A consistent theme from decades of developmental studies is that a relatively small group of morphogens generate variably sized concentration gradients that pattern tissues and organs into varying sizes and shapes. Yet, how such activities are regulated in diverse biological settings has remained a matter of active debate^1^. Specifically, whether some morphogens diffuse passively or rely on non-diffusive strategies to reach their targets has remained unclear (**Fig. 1A**)^2,3^. Deeper insight into the mechanisms of morphogen movement promises to unravel long-hidden biological mechanisms and could potentially inform a range of applications, including tissue engineering and drug delivery.

**Figure 1.**
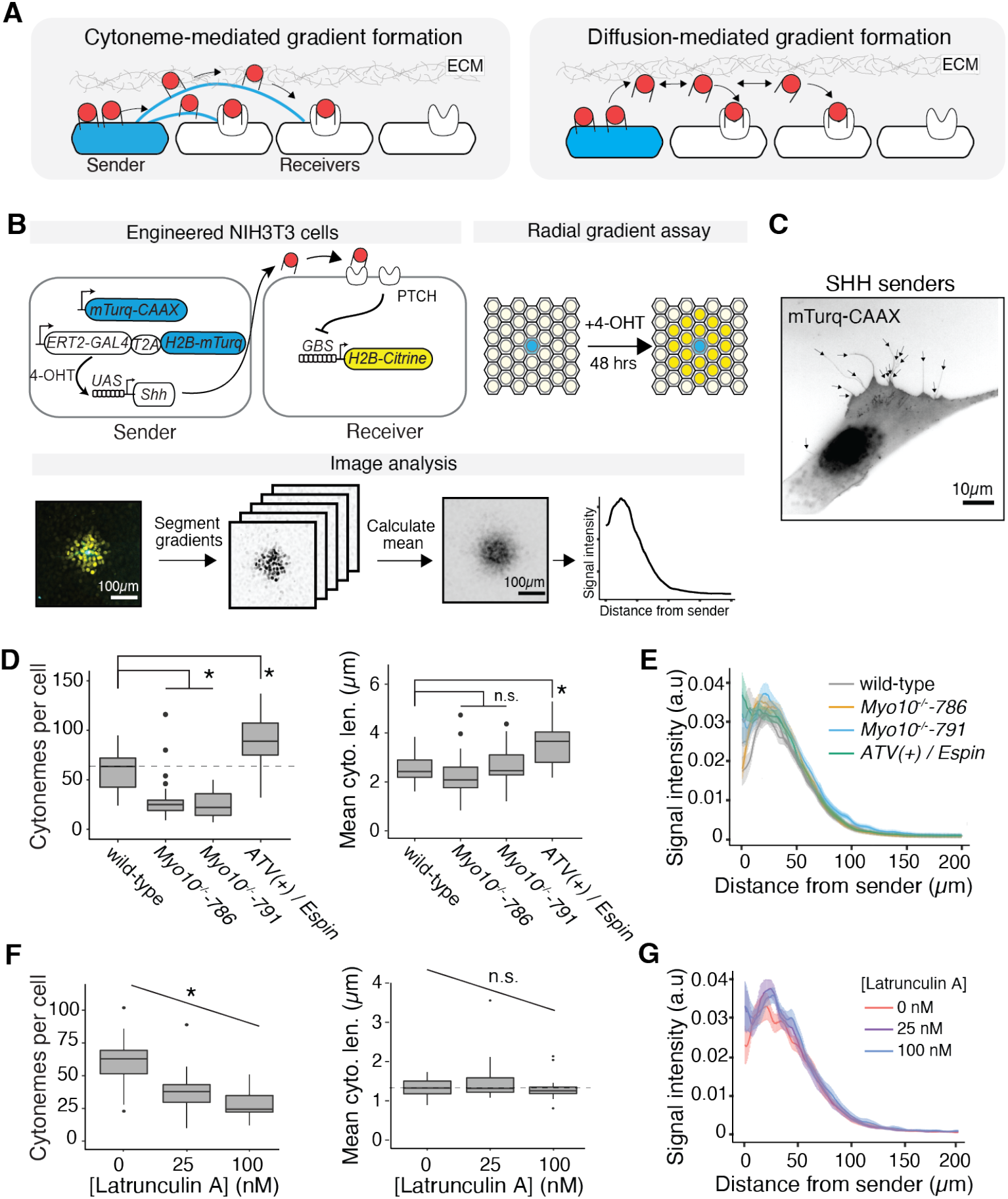
Cytoneme perturbations did not affect SHH signaling gradients. (**A)** Schematic of alternative models for morphogen transport. ECM reflects extracellular matrix. (**B)** Design and quantification of synthetic gradients. (**C)** A representative image of cytonemes on an NIH3T3 sender cell expressing mTurquoise2CAAX to label the cell membrane. Labelled sender cells were plated sparsely in a confluent lawn with unlabeled NIH3T3 cells. (**D)** Cytonemes were ablated in two independent *Myo10*^*(-/-)*^ clonal lines (denoted -786 and -791), and augmented in cell lines expressing the artificial transport vehicle tethering Espin to the tip of cytonemes (ATV(+)/Espin). Dashed line reflects the median of the parent NIH3T3 cell line, and (*) denotes statistically significant, using a two-tailed t-test and with Bonferroni-corrected threshold of a = 0.025. **(E)** Quantification of meta-gradients (i.e. average gradients) formed by sender cells with cytoneme deficiencies. Traces represent the mean fluorescence +/- the standard error of the mean (S.E.M.) for ∼30 gradients. (**F)** Latrunculin A limits actin polymerization, and caused cells to form ferwer cytonemes. Dashed line reflects the median of the parent NIH3T3 cell line without Latrunculin A, and (*) denotes statistically significant, using Pearson’s correlation test with a = 0.05. (n.s.) denotes not significant. (**G)** Quantification of signaling gradients formed in the presence of Latrunculin A.

Hedgehog (HH) family morphogens are at the forefront of this debate partly because of their lipophilic nature. Specifically, HH proteins have a palmitate on their N-terminus and cholesterol on their C-terminus, theoretically making them poorly soluble when secreted into the aqueous extracellular space^4,5^. Nevertheless, HH molecules can form patterning gradients that range from ∼25μm to ∼300μm^6–8^. To reconcile this contradiction, it has been suggested that HH is transported within specialized membrane protrusions called cytonemes, where HH can travel without contacting the aqueous extracellular environment (**Fig. 1A**)^9,10^. Indeed, in Drosophila wing discs, cytoneme length is positively correlated with HH signaling range^11^. Furthermore, chick limb bud fibroblasts secrete Sonic Hedgehog (SHH), and are known to form cytonemes that can transport SHH-eGFP^12^. Similarly, sparsely cultured mouse fibroblast sender cells can deliver SHH-mCherry to receiver cells via cytonemes^10^. In contrast, it is also clear that cultured SHH sender cells can secrete signaling-competent soluble SHH directly into the culture medium, suggesting that SHH signaling does not strictly require direct cell-cell contact^13–15^. The lack of clarity regarding how HH spreads makes it even more puzzling how the signaling range of HH can be tuned in diverse tissues.

Understanding how different transport mechanisms contribute to morphogen gradient formation requires observing and perturbing SHH delivery mechanisms in a context where the gradient can be quantitatively measured. Achieving this goal has been challenging in live embryos, because morphogens exist at low concentrations and access to highly sensitive microscopy is limited. To understand how SHH gradient formation is regulated, we previously reconstituted SHH signaling gradients in cultured mouse fibroblasts^16^. Here, using single-molecule imaging, we found that SHH diffused extracellularly and frequently transitioned among multiple diffusing states, some of which are confined to cell membranes. Based on this observation, we developed a new model, in which long-range gradient formation required both local diffusion associated with the cell membrane and transient diffusion away from cell membranes to jump between cells. The topology of the tissue, defined by the geometry of physical boundaries between cells, determines the number of jumps a molecule needs to complete in order to reach a certain distance. This “topology-limited” diffusion model predicted novel parameters that modulate gradient size, including the transition rates between confined and free populations. In fact, the chordate-novel Hedgehog-binding protein SCUBE1 (Signal peptide, CUB domain and EGF like domain containing) catalytically accelerated transitions between confined and free populations, suggesting that SCUBE1 expanded Hedgehog gradients at least partly by modulating its ability to cross diffusion barriers. Our results revealed novel and surprising mechanisms that evolution can leverage to shape morphogen gradients in a tissue- or species-specific manner.

## Results

### Hedgehog signaling gradients did not require cytonemes

To understand whether SHH was capable of extracellular diffusion, we first asked whether cytonemes on SHH-producing (“sender”) cells are required for delivering SHH from sender cells to responding (“receiver”) cells. To make precise measurements of SHH signaling gradients, we previously established an SHH signaling gradient assay, in which NIH3T3 mouse embryonic fibroblast cells were engineered to become SHH senders and receivers, and formed radial signaling gradients in co-culture (**Fig. 1B**).

The receiver cells express a synthetic SHH reporter driven by a consensus Gli binding site (GBS) array. We confirmed that upon SHH stimulation, the receiver cells expressed canonical SHH targets *Ptch1* and *Gli1*, and did not exhibit positive feedback on Hedgehog ligand gene expression, suggesting that the only source of Hedgehog ligand is the engineered sender cells (**Fig. S1**). We expressed a membrane-anchored mTurquoise2 to label the sender cell membrane, and found that NIH3T3 cells formed tube-like protrusions consistent with previous descriptions of cytonemes (**Fig. 1C**)^10,12^. We reasoned that if cytonemes are a major mode of SHH delivery, perturbations to cytoneme number or length in sender cells would have a quantitative impact on SHH signaling gradients. Thus, we generated two independent NIH3T3 SHH sender cell lines with genetic knockouts of *Myo10*, a non-canonical myosin known to localize to cytoneme tips and contribute to cytoneme extension (**Fig. S2**)^10,12,17^. Additionally, we developed a clonal SHH sender cell line expressing a Myosin XI-derived artificial transport vehicle (ATV(+)), which recruits the actin-bundling protein Espin to the tips of cytonemes, to stabilize long cytonemes^12^. Consistent with previous observations, we found that *Myo10*^*-/-*^ cells formed roughly half as many cytonemes compared to wild-type cells, and that cells expressing ATV(+)-Espin formed ∼50% more cytonemes compared to wild-type cells^10,12^ (**Fig. 1D**). Furthermore, whereas the cytoneme length distribution was unchanged in the *Myo10*^*-/-*^ cells, the median cytoneme in ATV(+)-Espin cells was ∼1.5-fold longer than in wild-type cells (**Fig. 1D**). None of the sender cell cytoneme perturbations affected the size of SHH signaling gradients, suggesting that sender cell cytoneme numbers and length did not control the size of SHH signaling gradients (**Fig. 1E**).

In addition to cytonemes on SHH-sending cells, cytonemes have been observed on SHH-receiving cells, and it has been suggested that receiver cells can “solicit” SHH from sender cells^18^. However, when we disrupted cytonemes on both sender and receiver cells with Latrunculin A, which limits actin polymerization by tracking actin in its monomeric state, SHH signaling gradients were not affected, despite a ∼50% reduction in the number of cytonemes^19^ (**Fig. 1F**,**G**). Finally, when we treated *Myo10*^*-/-*^ sender cells with Latrunculin A to further deplete cytonemes (∼75% reduction in cytoneme number compared to wild-type sender cells), we did not observe any defect in SHH signaling gradients (**Fig. S3**). Together, these results suggest that the size of SHH signaling gradients is not regulated by cytonemes.

### Hedgehog diffused extracellularly in four distinct populations

Next, we tested the alternative model in which SHH diffuses outside of cells and forms concentration gradients that determine the size of SHH signaling gradients. Using immunofluorescence analysis, we detected SHH on the sender cell membrane, but not in the extracellular space (**Fig. 2A**), consistent with prior observations that immunofluorescence often fails to detect extracellular morphogens likely because of their low abundance. Therefore, to visualize extracellular SHH we generated sender cells that secreted an SHH-Halo-Tag (SHH-Halo) fusion protein. SHH-Halo formed signaling gradients that were similar to those of wild-type SHH, suggesting that the HaloTag fusion preserves SHH activity (**Fig. S4**). We next labeled SHH-Halo with a cell impermeable fluorophore to detect extracellular SHH-Halo^20^. Using total internal reflection fluorescence (TIRF) microscopy, we observed strong signal on the sender cell membranes, and extracellular particles that formed a concentration gradient extending ∼50-100μm from the boundary of the nearest sender cell, consistent with extracellular diffusion of SHH-Halo (**Fig. 2A**,**B**). Notably, the size of the gradient measured by the extra-cellular SHH-Halo particles is slightly smaller than the gradient measured by the SHH signaling reporter (∼50μm vs ∼100μm). Two factors likely contribute to the small discrepancy. First, the SHH-Halo particle gradient was measured from the sender cell membrane to the SHH-Halo particle, whereas the signaling reporter gradient was measured from the sender cell nucleus to the receiver cell nucleus, using H2B-mTurquoise and H2B-mCitrine to measure sender and receiver positions respectively. Additionally, whereas TIRF imaging captured SHH-Halo at a discrete moment during gradient formation, live cell signaling reporters integrated the SHH concentration experienced during ∼48 hours of SHH signaling, and so are likely to be more sensitive to low amounts of SHH compared to TIRF-based imaging.

**Figure 2.**
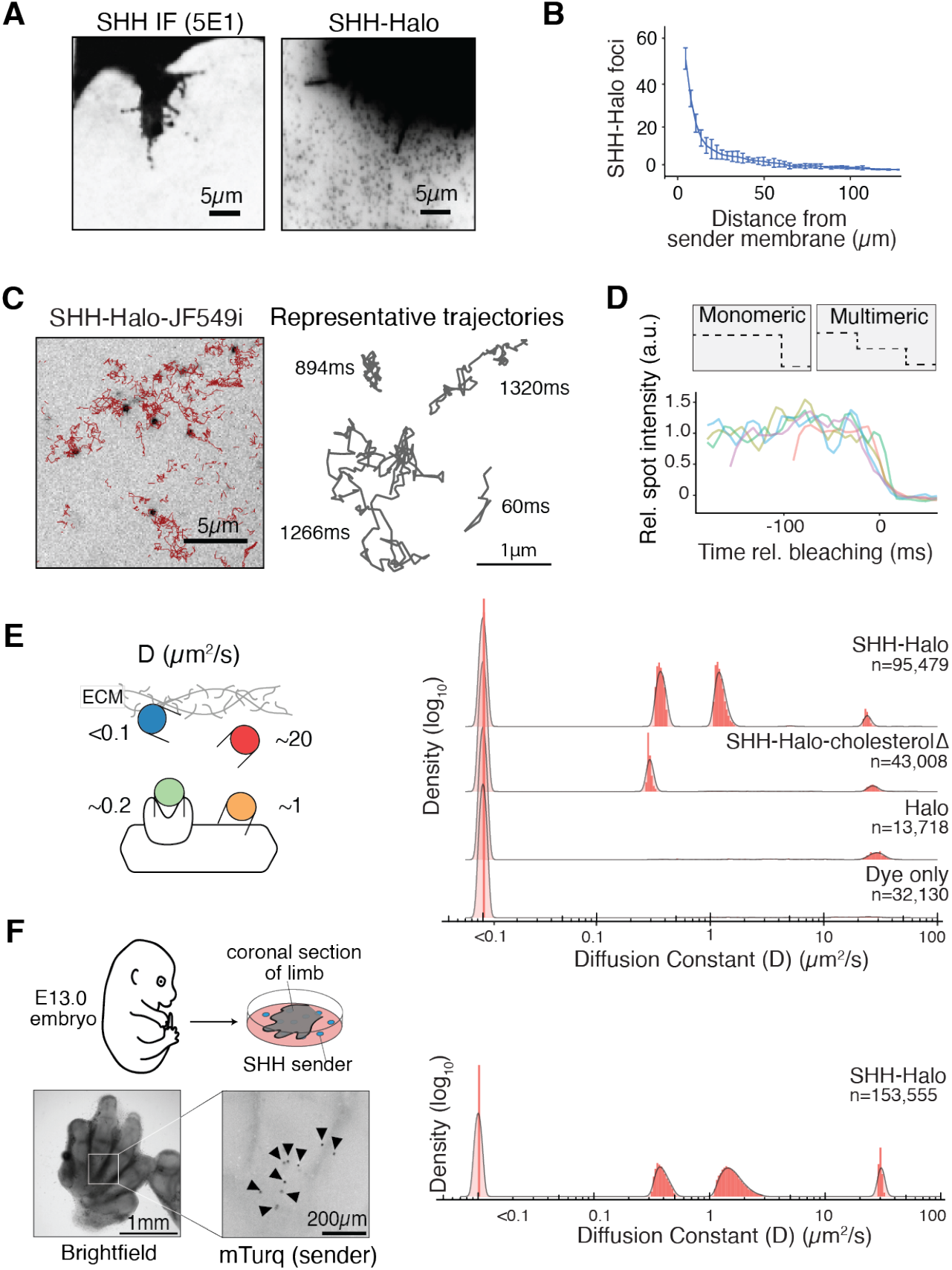
SHH diffused as heterogenous populations through the extracellular environment. (**A)** Microscopic detection of SHH, using immunofluorescence against native SHH (left) or imaging SHH-Halo with TIRF microscopy (right). Sender cells were cultured among a dense lawn of unlabeled wild-type NIH3T3 cells. (**B)** Quantification of SHH-Halo foci in A (right) as a function of distance from sender cell membrane, counted in 3μm bins. Each point reflects the mean number of particles in the given bin +/- S.E.M. for 8 sender cells. (**C)** (left) Timelapse imaging of SHH-Halo using TIRF microscopy. Red traces reflect single particle trajectories. (**D)** Particles of SHH-Halo bleached in a single step, suggesting that they were monodisperse. (**E)** Single particle diffusion rates reflected four discrete populations of SHH. Y-axis density values were log-transformed, and the Y scales were truncated for plotting. **(F)** E13.0 limb sections were cultured on top of sparsely plated NIH3T3 SHH-Halo sender cells, and single particle diffusion rates reflected four discrete populations of diffusive SHH-Halo.

To directly observe and quantify SHH diffusion, we used live imaging to track the movement of single particles of SHH-Halo during gradient formation^21^ (**Fig. 2C, Movie S1**). To ask whether each particle contained a single or multiple SHH molecules, we monitored fluorophore bleaching and found that single particles of SHH-Halo bleached in a single step, suggesting that each particle represented a single labeled SHH-Halo molecule (**Fig. 2D**). We localized SHH-Halo particles in frames captured every 6ms and calculated the maximum likelihood diffusion rate for each molecule in 18ms intervals: SHH-Halo diffused in four discrete populations with rates of ∼20μm^2^/s, ∼1μm^2^/s, ∼0.2μm^2^/s, and <0.1μm^2^/s (**Fig. 2E**). To test whether our microscopy conditions overlooked populations of molecules that were slower or faster than the observed molecules, we repeated the experiments using stroboscopic illumination over a four-fold slower acquisition rate (24ms exposure), or using continuous illumination over a two-fold faster (3ms exposure) acquisition rate and did not observe additional populations of SHH-Halo in the expanded sensitivity range (**Fig. S5**).

Given the drastic differences in diffusion rates between individual SHH molecules, we reasoned that each diffusing state likely reflected distinct interactions between SHH and the extracellular environment. To test whether diffusive populations reflected known features of SHH biochemistry, we took inspiration from prior experiments, in which SHH lacking its C-terminal cholesterol was found to form longer signaling gradients compared to wild-type SHH, presumably by altering SHH interactions in the extracellular space^22^. We generated an SHH-Halo allele that lacks cholesterol and found that the population of SHH-Halo molecules diffusing at ∼1μm^2^/s was dependent on the cholesterol modification, whereas other populations were unaffected (**Fig. 2E**). Interestingly, free lipids in a cell membrane are known to diffuse at ∼1μm^2^/s, suggesting that the observed SHH population diffusing at 1μm^2^/s could be anchored to the cell membrane by cholesterol^23^. Second, we found that secreted Halo alone diffused at ∼20μm^2^/s, similar to the fastest population of SHH-Halo (**Fig. 2E**). As an intracellular bacterial enzyme, Halo is not known to interact specifically with any mammalian protein, and the measured extracellular diffusion rate is strikingly similar to the measured intracellular diffusion rate of Halo inside mammalian cells^24^. Therefore, we reasoned that the population of Halo and SHH-Halo diffusing at ∼20μm^2^/s likely represents diffusion through the crowded extracellular environment without being confined by cell boundaries. Third, the population moving at 0.2μm^2^/s was absent in the Halo alone condition and likely reflected SHH-Halo specifically bound to membrane proteins including its receptor or coreceptors, because integral membrane proteins are known to diffuse at approximately ∼0.2μm^2^/s (**Fig. 2E**)^25^. The slowest population that diffused at <0.1μm^2^/s was likely associated with an insoluble factor in the extracellular environment, including either immobile extracellular matrix components or the glass cell culture substrate. Extremely slow molecules were present in experiments that tracked Halo alone and in the dye-only condition, suggesting nonspecific interactions between the Halo or the fluorescent dye and the cell culture environment could also contribute to the slowest population (**Fig. 2E**).

Next, we asked whether the molecular populations observed traveling through confluent NIH3T3 cell layers reflected how Hedgehog travels through a natural tissue. We cultured explanted E13.0 mouse limb tissue on top of sparsely plated SHH-Halo sender cells, and tracked SHH-Halo diffusing through the cultured primary limb tissue (**Fig. 2F**). In primary limb tissue, SHH-Halo diffused in four discrete populations with similar diffusion rates compared to SHH-Halo diffusing among NIH3T3 mouse embryonic fibroblasts, suggesting diffusion as discrete membrane-confined and membrane-unconfined populations is a natural feature of SHH (**Fig. 2F**).

### SCUBE1 expanded Hedgehog gradients

Morphogens can interact with diverse extracellular binding partners, and we asked whether such interactions can tune the SHH gradient size. SCUBE1 family proteins are chordate-specific secreted extra-cellular proteins that are required for normal Hedgehog-mediated patterning processes in vertebrates^26–28^. We expressed human SCUBE1 in NIH3T3 cells and generated conditioned media, which we added to the reconstituted SHH gradients. SCUBE1 expanded SHH signaling gradients by increasing both their amplitude and lengthscale in a concentration-dependent manner (**Fig. 3A**,**B**). Similarly, SCUBE1 expanded the SHH-Halo ligand and signaling gradients (**Fig. 3C, Fig. S4**). Importantly, we found that SCUBE1 did not alter the receiver cell response threshold to SHH, suggesting that SCUBE1 modulation of SHH gradients was not mediated by heightened sensitivity or by changes to the stability of SHH in the cell culture (**Fig. S6A**). Interestingly, mice and humans lacking SCUBE develop with shorter bones and with abnormal craniofac^i^ial structures compared to wild-type individuals, consistent with the model that SCUBE also expands Hedgehog gradients *in vivo*^29–31^.

**Figure 3.**
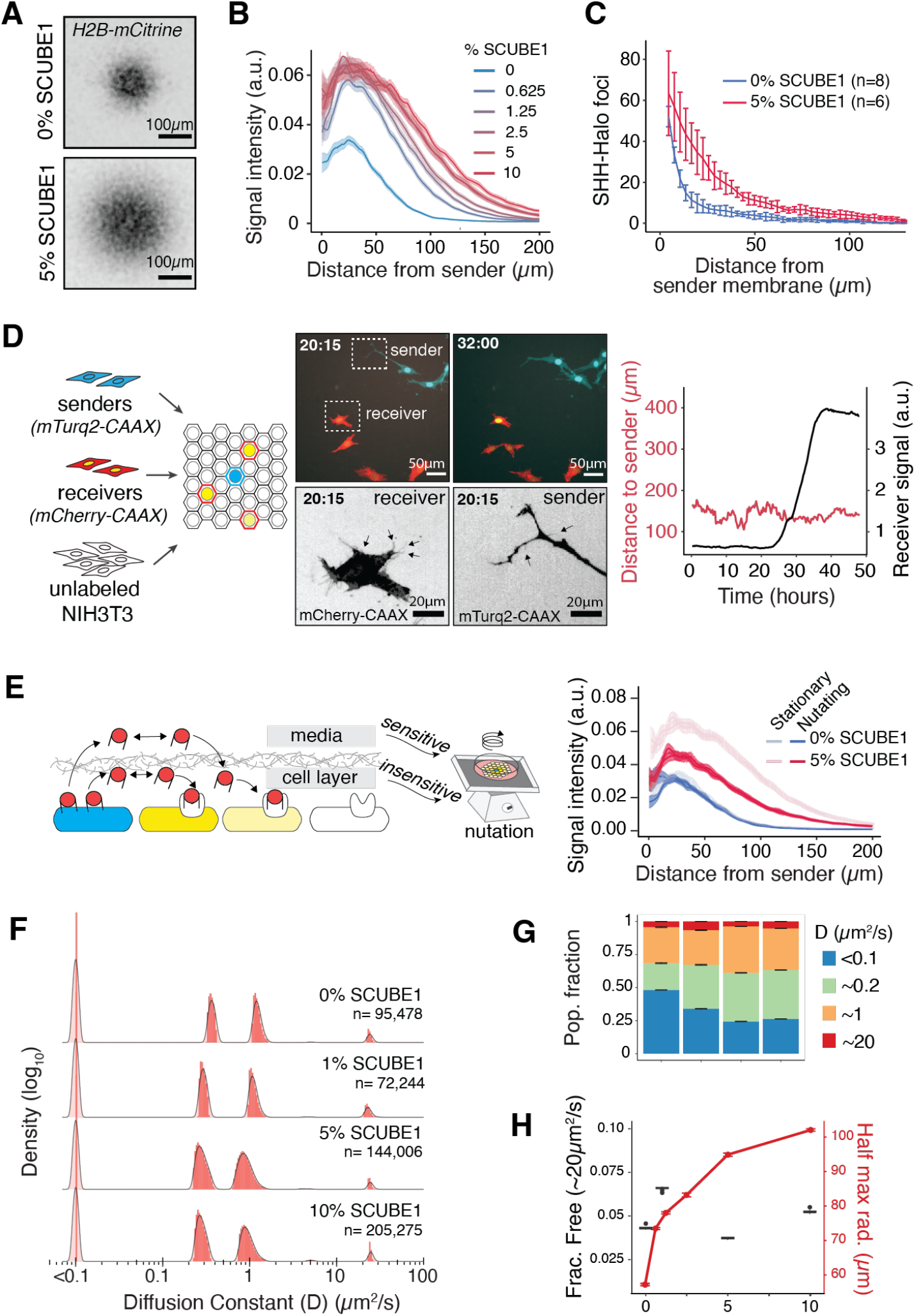
SCUBE1 expanded SHH gradients by modulating SHH movement. **(A)** Meta-gradients formed by single sender cells. (**B)** SCUBE1 expanded SHH signaling gradients in a dose-dependent manner. All cells were cultured in 10% conditioned media, mixed between wild-type conditioned media and SCUBE1 conditioned media to achieve the labelled %SCUBE1 conditioned media fraction. Trace reflects mean +/- S.E.M. (**C)** SCUBE1 extended the spatial distribution of SHH-Halo foci. SHH-Halo concentration gradient with 0% SCUBE1 is replotted from Fig. 2B. Trace reflects mean +/- S.E.M. (**D)** Sender, receiver and unlabeled bystander cells were co-plated, and senders were induced to signal. Still images are displayed from time-lapse imaging with 5% SCUBE1 conditioned media (Movie S2). Selected sender and receiver are displayed in grayscale to illustrate cytonemes. (**E)** In the presence of SCUBE1, SHH could travel either within the extracellular matrix of the cultured cells, or through the tissue culture medium. Nutation partially prevented gradient formation, only in the presence of SCUBE1. Trace reflects mean +/- S.E.M. (**F)** Histograms of diffusion coefficients for molecules observed across a range of SCUBE1 concentrations, measuring the diffusion rate over 3-interval (18ms) sub-trajectories. **(G)** Distribution of SHH among diffusive populations across a range of SCUBE1 concentrations. Error bars are +/- S.E.M. **(H)** Weighted average diffusion rate of SHH (black trace and axis) were not correlated to gradient lengthscale (red trace and axis) across a range of SCUBE1 concentrations. Error bars reflect mean +/- S.E.M..

Next, we asked whether the expansion of SHH gradients by SCUBE1 relied on diffusion or cytoneme-mediated sender-receiver contact. We genetically labeled sender and receiver cell membranes with mTurquoise2-CAAX and mCherry-CAAX respectively, and diluted the receivers with unlabeled NIH3T3 bystander cells to allow for better cell segmentation. Using timelapse imaging to track individual cells, we found that sender cells could transduce SHH signal to receiver cells in the presence of SCUBE1 despite never coming within ∼100μm of one another, suggesting that SHH diffused in the field of receiver cells (**Fig. 3D, Movie S2**). Importantly, SCUBE1 did not affect cytoneme properties (**Fig. S6B**). Together, these results suggest in the presence of SCUBE1, SHH gradients formed by diffusion.

In cultured SHH sender cells, SCUBE family proteins promote SHH release from SHH-producing cells into the culture media by interacting with the cholesterol moiety^13,14,32^. Therefore, one possibility is that SCUBE1 promoted SHH release into the tissue culture medium, and the secreted SHH could re-enter the cell layer and promote SHH signaling. Alternatively, SCUBE1 could promote SHH secretion or diffusion within the cell layer (**Fig. 3E**). To distinguish between these two models, we generated SHH signaling gradients on a standard laboratory shaker table to disrupt SHH movement through the tissue culture medium, and found that although SHH gradients were sensitive to nutation in the presence of SCUBE1, SCUBE1 still increased the amplitude and lengthscale of gradients formed under nutation (**Fig. 3E, Fig. S7**). This suggests that SCUBE1 promoted SHH mobility within the cell layer, in addition to promoting release of SHH into the tissue culture medium.

### Classic models of diffusion could not explain gradient expansion by SCUBE1

To understand how SCUBE1 expanded SHH gradients, we compared our results to the predictions of theoretical models of morphogen gradients. Under classic theoretical models, increasing the morphogen secretion rate increases the gradient amplitude, while increasing diffusion or decreasing degradation rates extends the gradient lengthscale^33^. Thus, the SCUBE1-dependent increase in signaling amplitude is likely caused by an increased amount of SHH released by sender cells. However, it is less clear how SCUBE1 modulated the gradient lengthscale. In cell culture, morphogens are degraded primarily through loss to the open tissue culture media or through receptor/coreceptor-mediated degradation. By growing cells under nutation, we found that SCUBE1 promoted SHH loss to the media (**Fig. 3E**), which is predicted to shrink SHH gradients. Additionally, we found that SCUBE1 did not affect signaling sensitivity, suggesting that it SCUBE1 is unlikely to affect receptor/coreceptor mediated degradation (**Fig. S6A**). Furthermore, if SCUBE1 physically protected SHH from an extracellular protease, cells should show a higher apparent sensitivity to SHH when treated with SCUBE1, however SCUBE1 did not affect the SHH response threshold, suggesting that SCUBE1 did not biochemically stabilize SHH. Although it is conceptually possible that SCUBE1 expanded gradients by preventing receptor-independent degradation, this mechanism is not supported in the literature and we considered it to be unlikely. Instead, we reasoned that by interacting with the cholesterol moiety on SHH^13,14^, SCUBE1 could influence how SHH interacts with the extracellular environment even after it is secreted from sender cells, and could therefore modulate its diffusion.

SHH diffused as heterogenous populations that associated with different extracellular environments, which matches the basic assumptions of the classic hindered diffusion model (**Fig. 2E**). Under this model, morphogens are thought to be either hindered (i.e. bound to a binding partner) or unhindered (i.e. freely diffusive), and the effective diffusion rate of a morphogen determined by the fraction of molecules that are freely diffusive^34^. Because SCUBE1 interacts with the cholesterol group on SHH to promote its release from sender cells^13,14^, and because the cholesterol moiety was required for SCUBE1-mediated Hedgehog gradient expansion (**Fig. S8**), we expected SCUBE1 to shift the SHH molecules from the membrane-bound fraction (∼1μm^2^/s) to the faster fraction (∼20μm^2^/s), in order to increase the effective diffusion rate.

To test our hypothesis, we performed single-molecule imaging to directly measure the diffusion rate of SHH. In the presence of SCUBE1, SHH-Halo still diffused as monomers in four states (**Fig. 3F, Fig. S9**). To our surprise, we only observed minor re-partitioning of molecules among the same four populations, and a large fraction of SHH still remained associated with the membrane, suggesting that SCUBE1 interacted transiently with cholesterol on SHH (**Fig. 3G**). Furthermore, despite the fact that SCUBE1 increased the gradient lengthscale, it did not increase the fraction of molecules in the freely diffusive population (**Fig. 3H**). This experimental finding was directly at odds with the central premise of the classic hindered diffusion model^1,34^.

### A topology-limited diffusion model uncovered novel ways to modulate gradient formation

To explain our experimental data, we reconsidered models of morphogen gradient formation. In classic morphogen gradient models, the extracellular space is always considered to be smooth and continuous. Here, we reasoned that the strong interaction between Hedgehog and the cell membrane could create distinct topological domains, such that Hedgehog molecules would either diffuse within the membrane (when associated with receptors or embedded in the cell membrane) or away from the cell membrane (when free or associated with the extracellular matrix). Under this model, the gaps between adjacent cells, even at submicron scale, would impose diffusion barriers for membrane-associated Hedgehog molecules. Consequently, Hedgehog molecules must move away from the cell membrane to overcome this barrier and form long-range gradients. Based on this logic, a tissue would function as a set of interrelated but discrete cells, with a set of cell-cell boundaries that form topological barriers and influence morphogen diffusion. Morphogens moving through spaces with discrete topological barriers must be governed by different design principles relative to those that describe the smooth and continuous spaces.

To test how topological barriers control Hedgehog diffusion, we implemented a model in which molecules diffused as two populations in a planar sheet of square cells but could transition between the two populations at specific rates. In the first scenario (the “topology-limited” model), one population was confined to a cell, and the other population could diffuse away from cells (and thus jump between cells). In the second scenario, both populations could jump between cells (equivalent to the classic hindered diffusion model) (**Fig. 4A**). We did not attempt to generate a specific model of mesenchymal cell arrangement, and instead used our physical model as an exploratory tool to understand whether topological confinement of morphogens resulted in non-obvious consequences for morphogen diffusion. When we simulated these scenarios, we identified parameters that could influence morphogen diffusion under the topology-limited diffusion model but not under the classic hindered diffusion model. Specifically, varying the transition rate between the two populations (without changing their relative abundance) was sufficient to regulate the effective diffusion rate and effective secretion rate of the molecules. This outcome suggests that a single biochemical parameter can regulate both SHH secretion and diffusion under the topology-limited diffusion model (**Fig. 4B, Fig. S10**). In contrast, under the classic hindered diffusion model, the effective diffusion rate was determined only by the steady state fraction of molecules in each diffusive population, and did not depend on the transition rate. This effect held true across all parameters tested, including across population fractions, cell sizes, and diffusion rates (**Fig. 4C,D, Fig. S11**).

**Figure 4.**
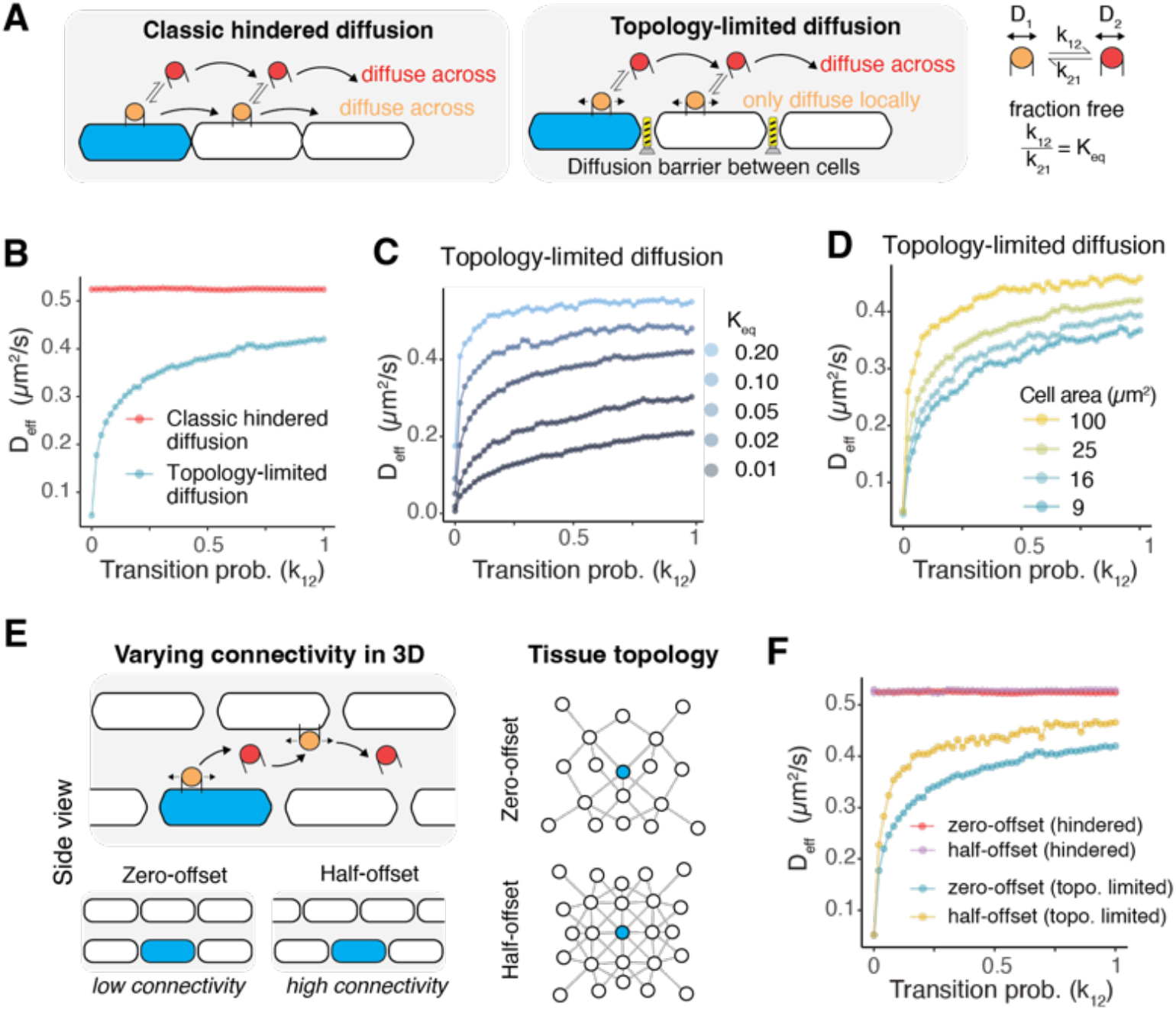
A topology-limited diffusion model predicted unexpected ways to modulate diffusion. **(A)** Schematic of classic hindered diffusion and topology-limited diffusion. Both models assume morphogens can transition between two diffusing populations. D_1_=0.5μm^2^/s and D_2_=1μm^2^/s are used for the subsequent simulation. **(B)** With the equilibrium fraction of each population fixed (K_eq_=0.05), the effective diffusion rate (D_eff_) is invariant to transition probability between the two populations under the classic hindered diffusion model, and sensitive to transition probability under the topology-limited diffusion model. The prediction for the topology-limited diffusion model holds across a range of K_eq_ **(C)** and cell area size **(D). (E-F)** Under the topology-limited diffusion model, distinct cell-cell arrangements give rise to topology-dependent effective diffusion rates. A higher connectivity (half-offset topology) leads to a larger D_eff_.

Given the rich complexity of networks of cell-cell contacts formed in cell culture and natural tissue, we reasoned that the extent of cell-cell contacts, which we call “tissue topology,” would be expected to influence gradient formation. Thus, we next simulated two planes of square cells over-growing one another (**Fig. 4E**). In one condition, we considered layers of cells in which each cell was perfectly stacked on top of another cell, such that each cell had four neighbors within its plane and one neighbor in the parallel plane. In an alternative condition, we offset the second layer of cells, such that each cell had four neighbors within its plane and four neighbors in the parallel plane, resulting in a greater number of topological neighbors compared to the condition in which cells are perfectly stacked (**Fig. 4E**). This design is conceptually equivalent to a three-dimensional simulation, where the third spatial dimension is small relative to the diffusion rate, as is likely true at the interface between two mesenchymal cells. Using this design, we found that cell-cell arrangement did not affect diffusion under the classic hindered diffusion model. However, under the topology-limited diffusion model, the effective diffusion rate shifted such that simulations with greater connectivity yielded faster effective diffusion rates (**Fig. 4F**). Importantly, although we perturbed tissue topology by considering alternative configurations of square cells, features like cytonemes or other membranous projections could similarly create topological “bridges” that could regulate the diffusion of morphogens if they significantly alter the cell-cell connectivity of the tissue in which the morphogen diffuses.

### SCUBE1 catalyzes interconversion of Hedgehog populations to promote gradient formation

Next, we asked whether SCUBE1 regulated the transition rates of Hedgehog between discrete diffusive populations by performing two related analyses based on our single-molecule imaging data. First, we recalculated the distribution of SHH-Halo diffusion rates across a range of integration time windows (**Fig. 5A**). Populations visible in a short integration window reflect the best approximation of the diffusion rate of pure populations of molecules, and populations that appear to travel with an intermediate diffusion coefficient only in long integration windows reflect molecules that transition between alternative populations during the integration interval^24,35–37^. We identified at least two populations that likely reflect molecules in transition between alternative molecular populations, including molecules in transition between ∼0.2μm^2^/s and ∼1μm^2^/s and between ∼1μm^2^/s and ∼20 μm^2^/s (**Fig. 5A**). Interestingly, both transitory populations emerged over shorter integration windows in the presence of SCUBE1, suggesting that each transition event occurred more frequently in the presence of SCUBE1 (**Fig. 5A**). Additionally, observation of molecules that transition between the “free” population and alternative SHH-specific biochemical populations suggests that molecules in all populations are structurally intact, and do not reflect a cleaved Halo tag or free fluorophore.

**Figure 5.**
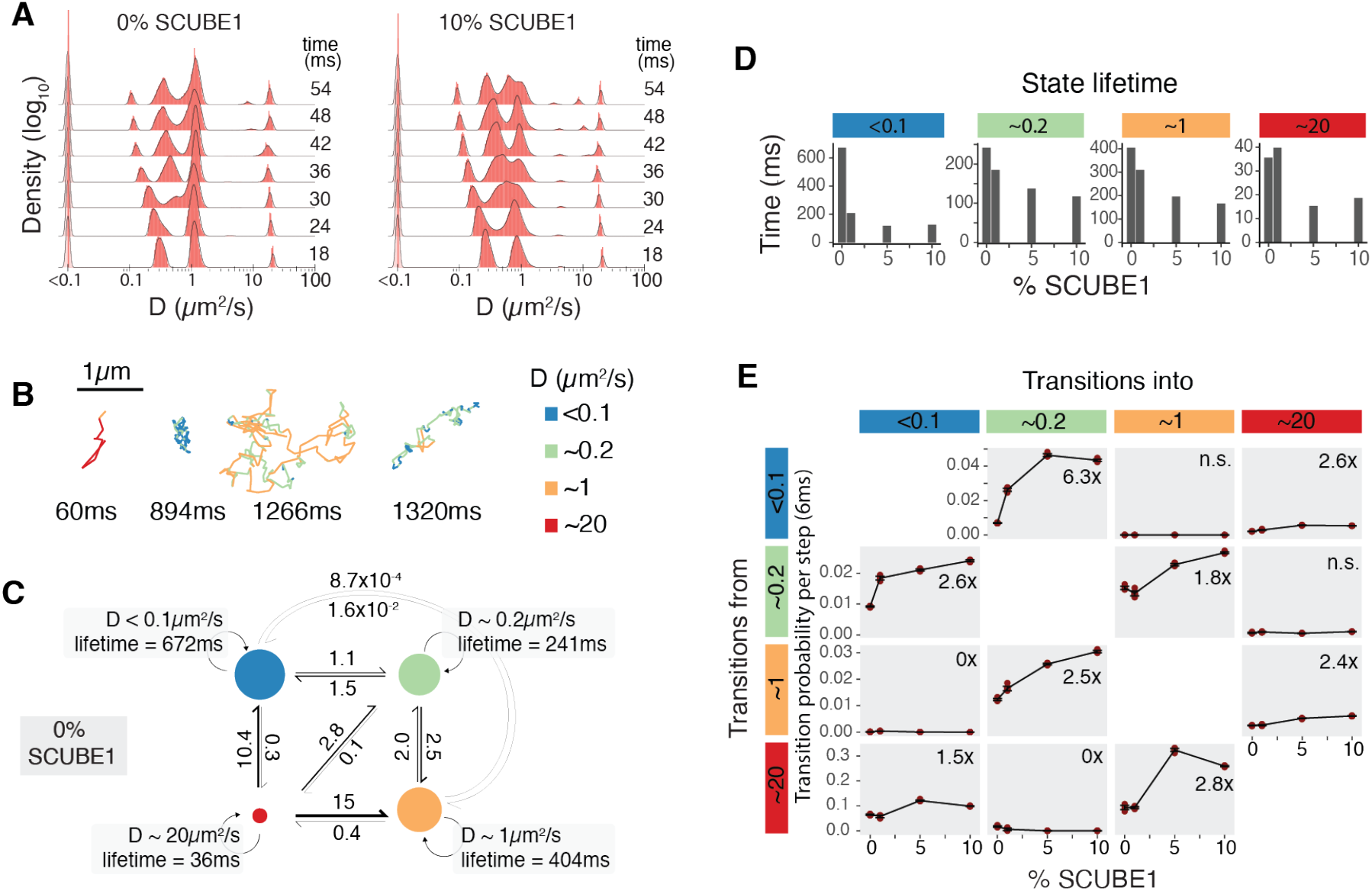
SCUBE1 catalyzes exchange between discrete diffusive populations of SHH. (**A)** Histograms of diffusion coefficients calculated over a range of integration windows. (**B)** *(top)* Representative single-molecule trajectories, color-coded by instantaneous diffusion rate. **(C)** State transitions diagram. The area of each circle represents the state occupancy. The number and weight of each arrow reflect the reaction rate (s^-1^) of each interconversion. (**D)** Lifetime of each state across SCUBE1 concentrations, estimated using a hidden Markov model. Estimate reflects the mean of 10 bootstrap replicates. **(E)** Transition probability per 6ms frame as a function of SCUBE1 concentration. Estimates from 10 bootstrap replicates plotted in red, and the mean +/- S.E.M. plotted as black lines. Numbers in the upper right of each panel reflect the fold-change in 10% SCUBE1 compared to 0% SCUBE1, for statistically significant changes, or n.s. where correlations were not significant (Pearson correlation with Bonferroni-corrected a = 0.00417).

In the second analysis, we used a recently published analysis method (ExTrack^37^) to jointly estimate the diffusion coefficients of four SHH-Halo populations as well as Markov parameters that described the interconversion rate between each population. We found that single trajectories of SHH-Halo transitioned between multiple diffusive populations (**Fig. 5B,C**). Furthermore, SCUBE1 acted as a catalyst to increase the transition rate between each SHH-Halo population and decrease the lifetime of each SHH-Halo population, without dramatically changing the net occupancy of each population (**Fig. 5D,E**). In order to transition between any pair of diffusible populations, we speculate that SHH molecules transit through a state in which the cholesterol group is transiently exposed to the aqueous environment. Such a transition presents a formidable free energy barrier, and we propose that SCUBE1 physically shields the cholesterol group to transiently decrease this free energy barrier. Such a mechanism would permit the rapid exchange between alternative populations of SHH-Halo without changing the steady-state occupancy of each fraction. Although SCUBE and SHH are known to form a biochemical complex^32^ we found no evidence that a stable freely diffusing SCUBE1-SHH complex existed during signaling (**Fig. 3F**). Therefore, we speculate that the SCUBE1-SHH transition state complex is distinct from the relatively stable state in which SHH diffused freely. Specifically, we speculate that when Hedgehog is in its freely mobile state, SHH palmitoyl and cholesterol moieties might interact *in cis* to exclude aqueous solvents, and that a function of SCUBE1 might be to help Hedgehog access this *cis*-interacting configuration. This mechanism is consistent with structural data that suggests that both hydrophobic moieties are attached to flexible linkers, and are spatially close enough to interact^38^.

## Discussion

By asking how SHH morphogen gradient size can be regulated, we discovered that prior models of morphogen diffusion could not adequately describe formation of SHH gradients^39–41^. Whereas prior models assume tissues to be uniform continuous spaces, our model considers tissues to be structured assemblies of discrete cells, separated by diffusion barriers. Under this model, molecules either diffuse in association with the cell membrane or are free to cross the diffusion barriers and jump from cell to cell. As a result, the transition rates between the confined and free populations determine the effective diffusion rate of a morphogen.

To develop this multiscale model of morphogen diffusion, we combined measurements of multicellular SHH gradients with new technologies for detecting and tracking single molecules of SHH in tissues. We found that SHH diffused as four discrete populations, two membrane-associated and two not associated with membrane, and that molecules rapidly transitioned between populations. Furthermore, the vertebrate-specific SHH-interacting protein SCUBE1 catalytically accelerated interconversion among the populations, and thus, increased the effective diffusion rate of SHH. SCUBE is known to promote SHH secretion^13,14^, however it was unknown if SCUBE could additionally affect SHH diffusion. Under the topology-limited diffusion model, secretion of SHH is similar to SHH diffusion, in that both processes can be mechanistically controlled by increasing the rate at which SHH transitions from the membrane-confined population to the free population that can jump between cells. Morphogen-interacting proteins exist broadly in most morphogen pathways, such as Chordin (for BMP), sFRP (for Wnt), and Glypicans (for multiple morphogens)^42–45^. It would be interesting to examine whether the catalytic activity of SCUBE1 is similar or different from the activity of other morphogen-interacting proteins. Interestingly, these knobs for tuning morphogen gradient size are often themselves regulated by the morphogen signal, forming feedback circuits that could ensure robustness of gradients to morphogen level fluctuations or scaling of gradients with tissue size^16,46–49^.

In addition to secretion and diffusion, natural signaling gradients can be shaped over longer timescales by mechanism like transcytosis, in which morphogens are internalized and then re-secreted, often with specific localization to promote requisite signaling outcomes^50^. Conceptually, mechanisms like transcytosis are similar to the topology-limited diffusion we observed during Hedgehog signaling, except that during transcytosis the confined populations of morphogens are intracellular rather than being membrane-associated. Importantly, however, topology would be a critical feature of pattern formation under either diffusive mechanisms or transcytosis-based mechanisms.

A prediction of our topology-limited diffusion model is that the degree of connectivity between cells can influence the formation of signaling gradients. However, we did not observe any differences in signaling gradient outcomes when we ablated or augmented cytonemes in mesenchymal cells, and corresponding experiments performed *in vivo* find incompletely penetrant lethality and minor patterning defects in viable animals^10,12,51^. We speculate that because mesenchymal cells are irregular and are tightly juxtaposed, cytonemes might represent a small fraction of the total extent of cell-cell contact, such that perturbations to cytonemes had minor consequence for the effective topology of the tissue. Notably, in epithelial sheets where the degree of intrinsic cell-cell contact is smaller, cytonemes might have a larger contribution to cell-cell connectivity, and therefore play a larger role in shaping the formation of morphogen gradients. Consistent with this observation, the most severe defects observed in mice with mutant cytonemes occur in the neural tube, which is comprised of epithelia cells that are known to form long-range cytonemes^10,51^.

The topology-limited model unified many ideas in the morphogen field that were previously considered to be unrelated or at odds. First, our model predicted that cell-cell connectivity could modulate the effective diffusion rate of SHH, suggesting that tissue structure is intimately related to morphogen gradient formation (**Fig. 4E,F**). Second, the model is generalizable to other morphogens and diffusible signaling ligands. In the case of SHH, membrane association is mediated by a combination of SHH-receptor interactions in addition to interactions between the SHH cholesterol moiety and cell membrane. For other morphogens without lipid modifications, cell confinement could be mediated by interactions with receptors, co-receptors, membrane-tethered matrix proteins or endocytic shuttling mechanisms, which could be regulated in an organism- or tissue-specific way to control both the fraction of signaling molecules that are confined and the transition rate between diffusive populations. Third, if the transition rate between populations is sufficiently fast, or if freely diffusive molecules are abundant, the topology-limited model is well-approximated by the classic hindered diffusion model. Therefore, the topology-limited diffusion model represents a general framework for understanding the basis of morphogen diffusion, and unifies morphogen gradient models that were previously considered to be incompatible. Taken together, our findings suggest that diffusion is a tightly regulated step in morphogen patterning, and point to a novel framework to understand how nature can modulate morphogen gradient sizes across evolution.

## Acknowledgements

We would like to acknowledge Ankur Jain, Luke Lavis, Zijian Zhang and Maria Barna for sharing critical reagents, Ankur Jain, Adam Martin, Peter Reddien and Mark Greenwood for advice on the manuscript, and Ankur Jain, Richard Hynes and Tom Kornberg for early advice and support, in addition to Brad Wierbowski, Julia Kim, Leah Wallach and the rest of the Li Lab for discussion and feedback through all stages of the project. This work was supported by National Institute of Health grants DP2HD108777 (PL), R00HD087532 (PL), DP2GM140938 (ASH), R33CA257878 (ASH), UM1HG011536 (ASH), 1K99GM151487-01 (GS), Allen Distinguished Investigator Award, a Paul G. Allen Frontiers Group advised grant of the Paul G. Allen Family Foundation (PL), National Science Foundation grant 2036037 (ASH), Jane Coffin Childs Fund Postdoctoral Fellowship (GS), and MathWorks Predoctoral Fellowship (MM).

## Author contributions

GS and PL conceptualized the project. GS performed the single-molecule tracking experiments and data analysis, with input and material support from DN and ASH. GS, MM and PL designed and performed cytoneme experiments. GS and PL developed the computational model. The manuscript was written by GS and PL, with input from ASH and MM.

## Data Sharing Plans

Plasmids used custom reagents used in this study are available upon request. Correspondence and requests for materials should be addressed to Pulin Li.

## Supplementary Information

Figures S1 to S11 Movies S1 to S2

## Materials & Methods

### Cells and cell culture

NIH3T3 mouse fibroblasts were used throughout the paper, and were cultured in DMEM + 10% Cosmic Calf Serum (GE Healthcare) supplemented with Penn/Strep/Glutamate and Sodium pyruvate. Imaging experiments were executed using DMEM without phenol red. Mouse limb explants were cultured in the same media, using 10% Tet-approved FBS (Takara) instead of calf serum. Sender and receiver cell lines were built as described previously^16^. Briefly, receiver cells were stably edited with Lipofectamine LTX and PiggyBAC transposase^52^ to express a fluorescent transcriptional reporter (P082 : *pGli(BS)-miniCMV-H2B-mCitrine*), and sender cells were stably edited to secrete SHH in response to stimulation with 4-OHT (P066 : *PGK-ERT2-Gal4-T2A-H2B-mTurquoise2;* P062 : *UAS-Shh*). Stable mTurquoise2-CAAX (JK054 : *pEF1a-mTurquoise2-CAAX*) and mCherry-CAAX (JK053 : *pEF1a-mCherry-CAAX*) cell lines were generated by editing established SHH sender or receiver cell lines with Lipofectamine LTX and PiggyBAC tranposase. The ATV(+)/Espin cell line was generated using Lipofectamine LTX and piggyBAC transposase based on a plasmid (pGS202 : *TRE-anti-GFPnb)-2A-Espin-sfGFP; EF1a-TetON3G*) described previously^12^ in established SHH sender cell lines. Cell lines for single molecule imaging were generated using Lipofectamine LTX and PiggyBAC transposase to insert Gal4-ERT2 and UAS-SHH-Halo, UAS-SHH-Halo-hintΔ (lacking cholesterol) or UAS-(SHH-sig-Peptide)-HALO in a single transfection (P066 : *PGK-ERT2-Gal4-T2A-H2B-mTurquoise2* ; pGS274 : *UAS-Shh-E130-HiBit-Halo-HA* ; pGS294 : *UAS-SHH-E130-HiBit-Halo-HA-hintΔ* ; pGS322 : *UAS-sigPep-HALO-HA*). All cell lines used in this study were clonally selected.

### Generating *Myo10* knockouts

SHH sender cells were transiently transfected with a Cas9-T2A-mCherry plasmid, encoding an sgRNA targeting the sequence AAGGATGGCTTGCTCGACGC. Transfected cells were sorted by FACS 24h after transfection for mCherry(+) cells, recovered for two days then sorted again to generate single-cell clones of *Myo10*^*-/-*^ cells. Clones were screened by PCR & Sanger sequencing, and knockouts were validated by immunoblotting.

### Measuring signaling gradients

SHH receiver cells were plated at ∼250,000 cells per square cm in 24-well imaging plates (i.e. ∼450,000 cells per well). Mitotic cells were counted by observing nuclear H2B-Citrine morphology, and cells were cultured until the culture stopped dividing. Next, sender cells were plated at a density of ∼100 cells per square cm (i.e. ∼200 cells per well) over-top the confluent receiver cells. Sender cells were induced with 250nM 4-OHT, and cultured for 48h before observing signaling gradients. For cells expressing TetA and (+)-end directed Espin, cells were cultured in 500ng/mL Doxycycline.

### SCUBE1 conditioned media

NIH3T3 cells were transfected with P100 (*pEF1a-Tet3G*) + P261 (*pTRE-miniCMV-hScube1*). Polyclonal cells were induced to secrete hSCUBE1 for 48 hours, and conditioned media was collected. Cell debris was removed by centrifugation. As a negative control, conditioned media was collected from wild-type NIH3T3 cells that were cultured under the same conditions. All experiments in which conditioned media was used included an equal fraction of conditioned media, mixing SCUBE1-containing conditioned media with wild-type conditioned media to achieve the stated SCUBE1 conditioned media fraction.

### Cytoneme counting

SHH sender cells with mTurquoise2-CAAX membrane labeling were plated at a density of ∼500 cells/cm^2^ in 24-well imaging plates (i.e. ∼1000 cells per well) and allowed to settle and attach to the substrate for ∼16 hours. After sender cells adhered, SHH receiver cells were plated over top at a density of ∼250,000 cells/cm^2^, and sender cells were induced to secrete SHH by adding 250nM 4-OHT. One day after induction, cells were imaged to count cytonemes. Microscopy was performed on a Nikon Ti2 with a 40x (NA 0.75) objective and a Zyla CMOS detector. Cells were imaged on 5 z-planes spanning 3.6μm, and cytonemes were counted manually in each plane using FIJI. Data were plotted using ggplot2 in R.

### Flow Cytometry

Receiver cells were cultured to confluence and then stimulated with dually lipidated recombinant SHH (R&D Systems, 8908-SH) either in the absence of presence of hSCUBE1 conditioned media, and analyzed 24 hours later using a BD Fortessa flow cytometer. Live, singlet cells were identified by gating for the modal population on forward scatter and side scatter, and identical gates were used for each experimental condition.

### Immunoblotting

Immunoblotting was performed using standard procedures. Briefly, ∼10^6^ cells were trypsinized ∼10m in 0.05% trypsin, then trypsin was neutralized with tissue culture medium, and cells were centrifuged for 5 minutes at 500xg. Cells were washed 2 times with PBS, then lysed on ice for 10 minutes in 100μL Tris-Triton lysis buffer (10mM Tris pH 7.4; 100mM NaCl; 1mM EDTA; 1mM EGTA; 1% Triton X-100; 10% glycerol; 0.1% SDS; 0.5% deoxycholate) with protease inhibitors (cOmplete Mini, Roche). Extracts were mixed with reducing agent (Life Technologies) and LDS sample buffer (Life Technologies), then heated at 95°C for 10 minutes. Extracts were chilled on ice, then centrifuged for 2 minutes at 15,000xg, and 10μL soluble protein was loaded into a pre-cast NuPAGE 3-8% Tris-Acetate gel (Invitrogen). Proteins were transferred using an iBlot2 (Invitrogen) at 25V for 10 min. The membrane was blocked with Licor TBS blocking buffer for 2 hours at room temperature, then detected overnight at 4°C with rabbit anti-Myo10 polyclonal primary antibody (PA5-55019, Invitrogen) at a 1:500 dilution. As a loading control, the membrane was labeled with mouse anti-fibronectin primary antibody (66042-1, Proteintech) at a 1:500 dilution. LiCor goat anti-mouse and goat anti-rabbit secondary antibodies were used to detect primary antibodies, and membranes were imaged using a Licor Odyssey Clx fluorescence scanner.

### Immunofluorescence

SHH sender and receiver cells were plated as for measuring SHH signaling gradients, and induced to signal for 48 hours. Cells were fixed for 30 minutes in 4% formaldehyde in PBS, and were permeabilized for 10 minutes at -20°C with a solution of equal parts methanol and acetone. Cells were blocked for 1 hour at 37°C in IP blocking buffer comprising 2.5% normal goat serum in PBS, with 0.1% Tween-20. Cultures were incubated overnight at 4°C with mouse anti-SHH primary antibody (cat #5E1, DSHB), diluted 1:50 in IP blocking buffer. Cells were washed 2x with PBS, and incubated for 4h with goat anti-mouse secondary antibody conjugated to AF-594 (Cat # A-11032, Invitrogen) diluted 1:2000 in IP blocking buffer. Cells were washed 3x in PBS and incubated at 4°C overnight prior to imaging.

### Halo fusion protein design and labelling

We inserted a cassette that encoded gly-HiBit-gly-ser-Halogly-ser-HA-gly in the mouse SHH coding sequence at position E130 (counting from the signal peptide). We chose this insertion site based on a previously published luciferase insertion at the same location that preserved the interaction between hSCUBE1 and SHH^53^.

To track single particles of SHH-Halo, SHH-Halo sender and wild-type NIH3T3 receiver cells were plated as for measuring SHH signaling gradients, except sender cells were plated at a much higher density (∼1% of final cell number). Cells were cultured in 35mm MATEK glass bottom dishes (uncoated). Cell culture media were supplemented with conditioned media as required for the experiment and induced to signal for ∼18 hours. For experiments utilizing hSCUBE1 conditioned media, the total amount of conditioned media was held constant between conditions by supplementing tissue culture medium with conditioned media from wild-type NIH3T3 cells. After ∼18 hours, Halo-ligands (JF549i or JF635i, gifts from Luke Lavis) were added to a final concentration of 50nM, and incubated at room temperature for 5 minutes. Cells were gently washed 3 times with pre-warmed tissue culture medium, and all washes utilized the same media stock as the original experiment such that the concentration of hSCUBE1 in each well did not change during the wash steps.

### Single-molecule imaging

Single-molecule concentration gradient experiments were conducted using a 505 Dragonfly TIRF microscope (Andor Technologies), with an iXon Ultra 888 EMCCD using a 60x TIRF objective (NA 1.49). Cells were imaged in a stage-top humidified incubator (OKO labs) at 37°C and 5% CO_2_. Data were analyzed using custom scripts calling Fiji and Track-mate^54–56^. Particles were counted in 3μm radial bins, and smoothed over 9μm intervals for plotting.

Single-molecule tracking experiments were conducted on a custom-built Nikon (Nikon Instruments Inc., Melville, NY) Ti-2E microscope equipped with a 100x (NA 1.49) oil-immersion TIRF objective (Nikon apochromat CFI SR HP Apo TIRF 100x Oil), sCMOS camera (Teledyne Photometrics, Tucson, AZ, Prime 95B), a perfect focus system to correct for axial drift and motorized dual galvo laser illumination system (Gataca Systems, Massy, France, iLas2), which allows an incident angle adjustment to achieve TIRF or highly inclined and laminated optical sheet illumination^57^. The incubation chamber maintained a humidified 37°C atmosphere with 5% CO_2_ and the objective was similarly heated to 37°C for live-cell experiments. Excitation was achieved using a 561 nm laser line (1 W, Genesis Coherent, Santa Clara, CA) for JF549i. The excitation lasers were modulated by an acousto-optic Tunable Filter (AA Opto-Electronic, France, AOTFnC-400.650-TN) and triggered with the camera TTL exposure output signal. The laser light is coupled into the microscope by an optical fiber and then reflected using a multi-band dichroic (405 nm/488 nm/561 nm/633 nm quad-band, Semrock, Rochester, NY) and then focused in the back focal plane of the objective. Fluorescence emission light was filtered using a single band-pass filter placed in front of the camera using the Semrock 593/40 nm bandpass filter. The microscope, cameras, and hardware were controlled through the NIS-Elements software (Nikon), and data were analyzed using Fiji, Trackmate, SaSpt and ExTrack as required for the experiment^24,37,54–56^.

For single-particle tracking with exposure times of 3ms and 6ms, frames were captured under continuous illumination. For tracking with exposure time of 24ms, the main excitation laser 561nm was pulsed for 6ms using the acousto-optic tunable filter, to achieve 6ms excitation times during a 24ms frame capture^21^. For 3ms and 6ms exposures, the ROI was constrained to achieve the necessary data transfer speeds, by capturing ROIs with y dimension 146px and 290px respectively (ROI y-dimension is limiting for data transfer speeds from CMOS detectors). The laser angle was chosen to give TIRF or very-near TIRF excitation, with an estimated excitation depth of ∼100nm measured from the top of the cover glass surface.

### Limb explant sectioning

Surplus forelimbs and hindlimbs from E13.0 embryos were dissected and embedded in 4% low-melt agarose, then sectioned on ice using a Leica VT1200S vibratome with a step size of 100μm. After sectioning, individual sections were removed from the agarose and plated on MATEK imaging plates pre-seeded sparsely with SHH-Halo sender cells. Explants were cultured with a minimal amount of media overnight to promote adhesion of the glass substrate. After overnight culture, limb sections adhered to the culture dish and formed a primary cell-derived sheet, with sparsely plated SHH-Halo sender cells incorporated into the cell layer. Sender cells were induced to secrete SHH-Halo for 16-24 hours before imaging SHH-Halo. Media was identical to that used for NIH3T3 cells, except 10% tet-approved fetal bovine serum (Takara) was used in lieu of cosmic calf serum. Whole explants were imaged using an inverted epifluorescence microscope with a 10x objective, then moved to a TIRF microscope for single molecule imaging experiments. Embryos were surplus from experiments approved under mouse protocol 0920-101-23, overseen by the MIT committee on Animal Care.

### Bleaching analysis

The mean fluorescence of each spot was normalized such that the mean pre-bleached intensity was set to 1, and the mean post-bleached intensity was set to 0, and the fluorescence traces were smoothed with a rolling mean over a 3-frame window for plotting. To plot timepoints after the end of a trajectory, we measured the fluorescence at the site of the particle’s last detected position. Representative trajectories were manually identified, and reflect a range of diffusive populations.

### Single-Particle Tracking (SPT) data analysis

Particles were detected using custom scripts to call Trackmate^56^. Particles were assumed to have a diameter of 600nm, quality score of 3 or 3.5 for 6ms or 24ms imaging intervals (based on manual QC) or 2.5 for 3ms imaging intervals, and were connected into trajectories with a maximum linking distance of 1.71μm. Discontinuous trajectories were not allowed.

We used two methods to estimate the diffusion rates and localization errors for SPT data. First, we used the saSPT python package to fit a state array representing the posterior probability of each trajectory under a family of error and diffusion rates^24^. We performed saSPT allowing localization error between 0.025μm and 0.05μm, and sampled diffusion rates from 0.01μm^2^/s to 100μm^2^/s, with integration windows set for each experiment as described in the main text. For each sub-trajectory, we marginalized over error rates and calculated the posterior maximum likelihood diffusion rate, then displayed histograms of the maximum likelihood diffusion rates using ggplot2 in R^58^. The apparent maximum likelihood diffusion coefficient calculated by saSPT depended on the integration interval. Because SHH-Halo transitioned between diffusible populations very rapidly, we were forced to consider extremely short integration times (3 localization intervals, 18ms) introducing error in the measurement of each population’s diffusion coefficient. Therefore, we also measured diffusion rates using the python package ExTrack, which was developed to fit a hidden Markov model to single particle tracking data to jointly estimate diffusion rate, state occupancy and transition probability^37^.

To display diffusion rate data on raw trajectories (**Fig. 5B**), we generated 4-interval rolling windows around each localization, and used SaSPT to calculate the maximum likelihood diffusion coefficient in each rolling window. We colored each displacement by the implied local diffusion coefficient using -2:+2 localizations to calculate the instantaneous implied local coefficient.

### HMM analysis

We used the python implementation of ExTrack to fit a hidden Markov model to the observed single particle tracking data^37^. We allowed ExTrack to consider 4-interval sub-trajectories drawn from trajectories that included 3-100 detections, and to fit parameters representing 4 populations of molecular states. We used ExTrack to jointly estimate the diffusion rate of each population, the occupancy of each population and transition rate between each population under the assumption that transitions occur at most once per 6ms observation interval. To estimate confidence intervals, we performed 10-fold delete-10% jackknife sampling on our data based on a common set of priors, which we generated by fitting the ExTrack HMM model to 50% the same underlying data with conservative priors. The results of ExTrack’s HMM fit were plotted in R using ggplot2^58^. We noted that population fraction implied by the transition matrix was slightly different than the observed instantaneous population fraction estimated by ExTrack or SaSpt, which reflects the fact that ExTrack did not converge to a global maximum likelihood estimation for every transition rate. As a practical matter, observing transitions into or out of the fastest population is considerably harder than measuring the population frequency of the fastest population, because fast-moving trajectories often leave the focal plane before they experience a transition to a different diffusible population. When interpreting the transition matrices, we assumed that the bias towards slower-moving molecules affected each experimental condition equally.

### Agent-based simulations

Numerical simulations of SHH diffusion were performed in MATLAB, and results were visualized using ggplot2 in R^58^. We initialized a 2D lattice of square cells (5μm x 5μm), and randomly initialized SHH molecules (“agents”) at a random position in a “sender” cell. Each agent was assigned to a population randomly, such that 5% of the molecules were assigned to the “free” fraction and 95% to the “constrained” fraction. Agents were allowed to diffuse by choosing their x and y displacements at each time step independently from a normal distribution described by 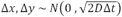 where Δt was 6ms and D was the diffusion coefficient of the corresponding population. Constrained molecules were assigned a diffusion rate of 0.5μm^2^/s and free molecules were assigned a diffusion rate of 1μm^2^/s. At each time step, a molecule was allowed to transition between populations, and the transition rate into the constrained population was assumed to be 20-fold greater than the corresponding transition rate into the free population, such that the population of molecules contained on average 5% molecules in the free state. For the “classic hindered diffusion” models, molecules were allowed to diffuse across cell boundaries without impediment, however for “topology-limited” diffusion models, molecules in the constrained population were prevented from jumping across cell boundaries and experienced a reflective boundary at the cell border. Only the molecules in free population were allowed to jump across cell boundaries.

We used the agent-based model in two ways. First, to estimate the effective diffusion rate (D_eff_), we initialized 5,000 agents in a single sender cell and allowed them to diffuse through the 2D lattice. At each time step, we measured the mean squared displacement of molecules from the center of the sender cell, and performed the simulation until the slope (mean square displacement (MSD) vs time) reached steady state (120s of simulated time, 20,000 time steps). To measure the implied D_eff,_ we calculated the slope of MSD over time at steady state and divided by 4, based on the relationship MSD(t) = 4Dt for diffusion in 2 dimensions.

For bilayer simulations, we assumed that molecules diffused in a differential volume at the interface between two cell layers, which the two layers of cells were either perfectly stacked or offset by 50% on both dimensions. When molecules transitioned from the free to the confined population, they randomly chose a cell based on their position, such that 50% of molecules entered the “upper” cell layer, and 50% entered the “lower” cell layer. Network diagrams were plotted in R using the library iGraph.

To estimate the effective flux (J_eff_), we performed similar simulations except we set the number of SHH molecules within the sender cell to be constant, with 2000 molecules at each time step. This assumption is equivalent to fixing the concentration of the morphogen source at a constant value, and comes from our intuition that the sender cell membrane is saturated with morphogens during cell signaling. If a molecule left the sender cell, it was added to a table of secreted molecules. At each time step, the secreted molecules were also allowed to diffuse (exactly as described in D_eff_ estimates), and molecules that returned to the sender cell were removed from the simulation. The total number of molecules in the receiver cell field was recorded at each time step. The simulation continued until the flux reached a constant rate (40,000 time steps representing 240s of simulated time), and the effective flux (J_eff_) was calculated as the steady state slope of the number of secreted molecules as a function of time under each molecular transition rate.

