## Supplemental Information for "Diffusion barriers imposed by tissue topology shape morphogen gradients"

**Supplementary Information**

Figures S1 to S10

Movies S1 to S2

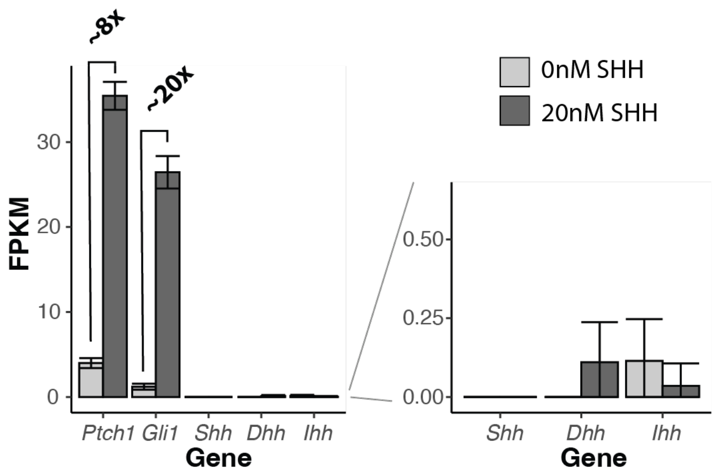


**Figure S1. RNA sequencing of cells induced with recombinant SHH**

Canonical SHH target genes *Ptch1* and *Gli1* are induced by stimulation with SHH, but Hedgehog paralogs *Shh*, *Dhh* and *Ihh* were not induced by recombinant SHH. Data re-analyzed from Li *et al.* [16]

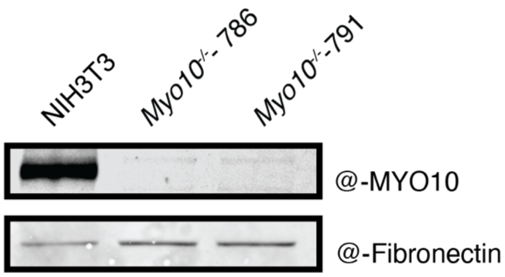


**Figure S2. Validation of *Myo10* knockout by immunoblotting**

*Myo10*$^{-/-}$ mutants produced no detectable MYO10 protein, using total cell extract.

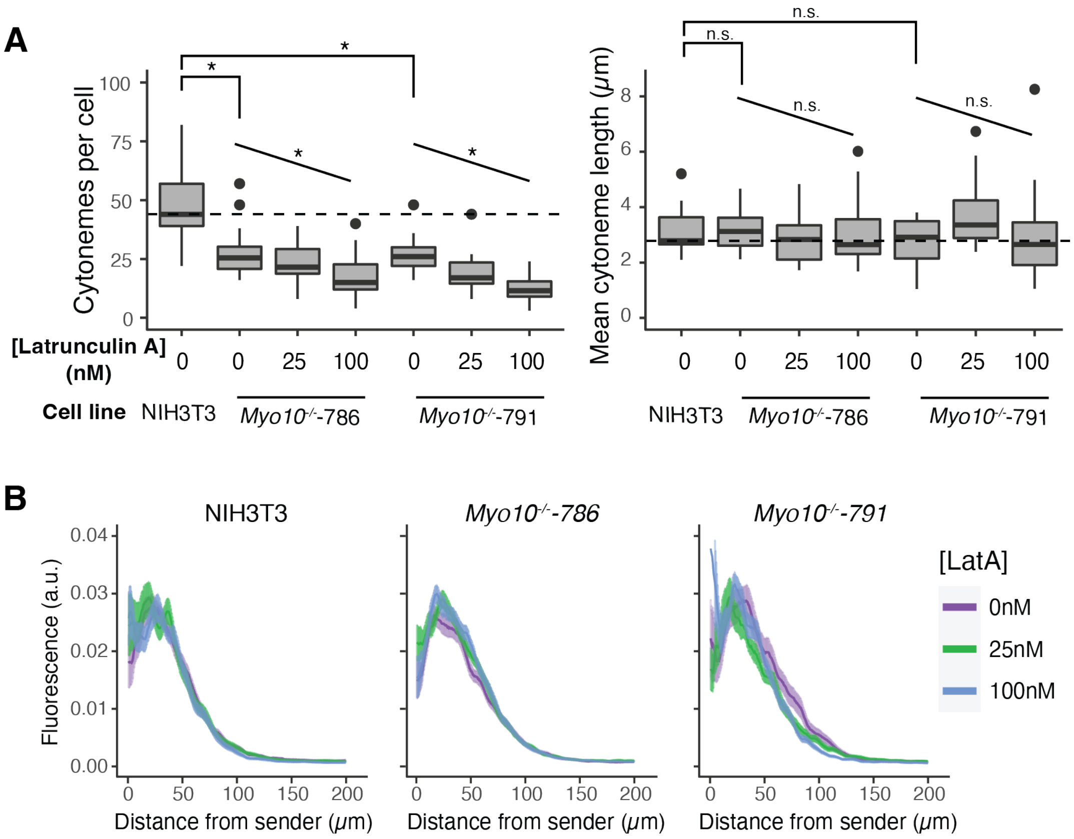


**Figure S3.** ***Myo10$^{-/-}$*** **cells cultured in the presence of Latrunculin A did not have signaling gradient defects**

**A.** Latrunculin A interacted additively with *Myo10$^{-/-}$*. (left) Chemically ablated mutant cells had fewer cytonemes compared to the singly-perturbed cells. Untreated *Myo10$^{-/-}$* cells had fewer cytonemes compared to wild-type cells (* denotes statistically significant t-test with Bonferroni-corrected $\alpha = 0.0125$), and treated cells further decreased cytoneme number (* denotes statistically significant Pearson correlation, with Bonferroni-corrected $\alpha = 0.0125$). (right) Latrunculin A did not change the length distribution of cytonemes. Untreated *Myo10$^{-/-}$* cells formed cytonemes that were similarly long compared to wildtype cells (n.s. denotes not statistically significant, t-test with uncorrected $\alpha = 0.05$), and latrunculin did not quantitatively shorten cytonemes (n.s. denotes not statistically significant Pearson correlation, with uncorrected $\alpha = 0.05$). **B.** SHH signaling gradients were not affected by Latrunculin A in *Myo10$^{-/-}$* cells.

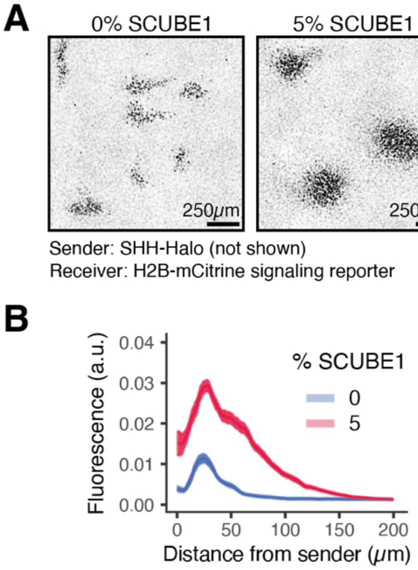


**Figure S4. SCUBE1 extended signaling gradients formed by SHH-Halo, related to Figure 3.**

**A.** Representative fields from co-cultured SHH-Halo sender cells with wild-type receiver cells. **B.** Quantification of meta gradients summarized in (A).

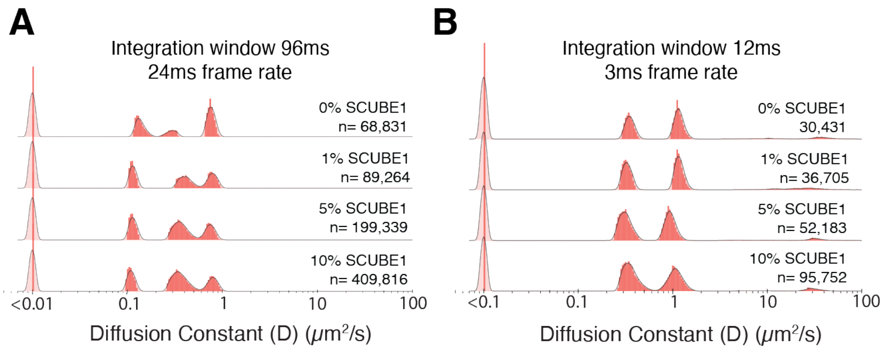


**Figure S5. Validating the temporal resolution of the single-molecule imaging**

Imaging SHH-Halo diffusion with an expanded range of imaging parameters did not reveal any population of SHH-Halo that was not captured with a 6ms frame rate. **A.** 24ms imaging frequency captured an intermediate diffusion population consistent with transition between the adjacent diffusion populations (discussed at length in Fig. 5), but did not identify any intermediate populations between 0.01μm$^2$/s and ~0.2μm$^2$/s. **B.** 3ms imaging frequency did not detect any molecules moving faster than the population at ~22μm$^2$/s that we identified using 6ms imaging frequency, although it is possible there are molecules moving at a rate faster than 50μm$^2$/s, which we would not be able to observe due to technical limitations.

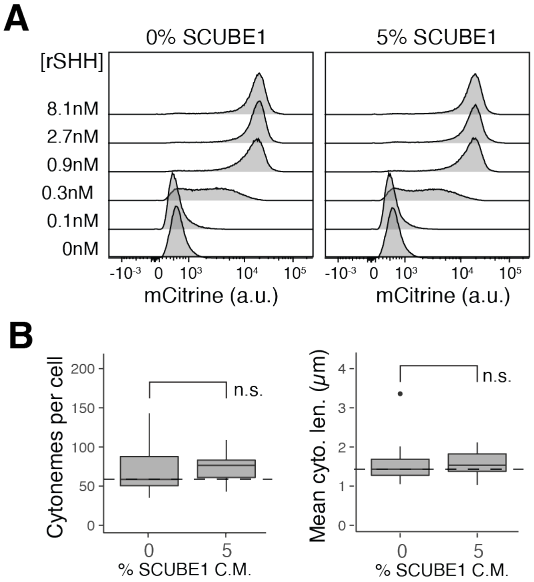


**Figure S6. SCUBE1 did not affect receiver cells' sensitivity to SHH or NIH3T3 cell cytonemes**

**A.** Receiver cells were cultured either in the absence or in the presence of 5% SCUBE1 conditioned media (C.M.), and stimulated with recombinant SHH protein. Receiver cells responded to the same dose of SHH regardless whether they were cultured in the absence or presence of SCUBE1conditioned media. Each sample used identical gates on forward and side scatter to identify monodisperse, healthy cells, and each histogram reflects 100,00 healthy cells. **B.** SCUBE1 did not affect cytoneme number or length in sender cells.

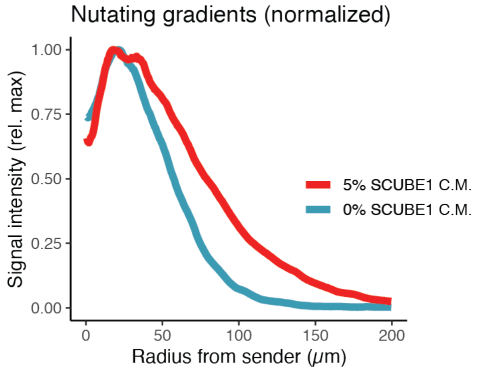


**Figure S7. SCUBE1 extended the lengthscale of SHH signaling gradients under nutation**

Gradients were scaled to their maximum value to observe the relative lengthscale in the presence of SCUBE1. Underlying data is identical to Fig. 3E.

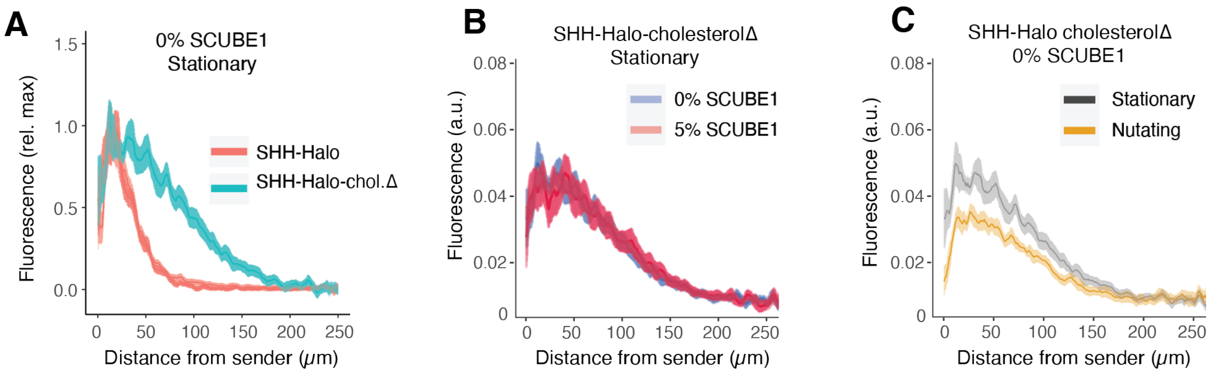


**Figure S8. SCUBE1-mediated gradient expansion was related to the SHH cholesterol modification**

**A.** SHH-Halo-cholesterolΔ formed longer signaling gradients compared to fully modified SHH-Halo. Trace reflects mean +/- S.E.M. for >10 gradients. **B.** SHH-Halo-cholesterolΔ gradients could not be expanded by SCUBE1. Trace reflects mean +/- S.E.M. for 10 gradients. **C.** SHH-Halo-cholesterolΔ gradients were partially sensitive to nutation. Trace reflects mean +/- S.E.M. for >9 gradients.

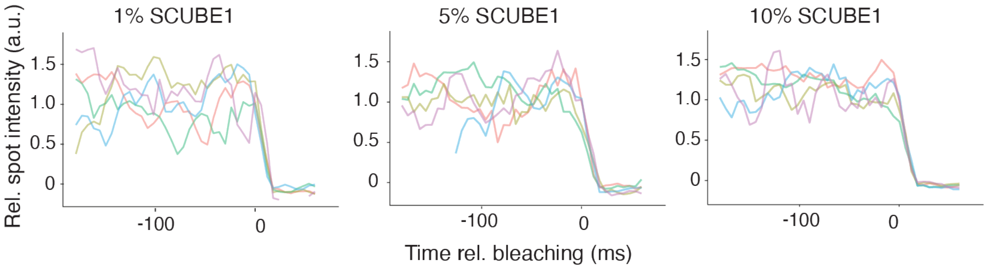


**Figure S9. Bleaching analysis of SHH-Halo particles in the presence of SCUBE1**

Particles of SHH-Halo did not show evidence of multi-step bleaching in the presence of SCUBE1, suggesting that particles were monodisperse.

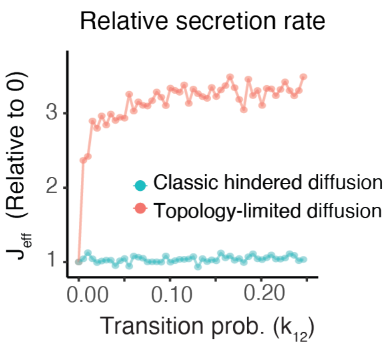


**Figure S10. Agent-based simulation of effective secretion rate (flux) across a range of transition rates**

Under the topology-limited model, increasing the transition rate between constrained or free populations increased the effective secretion rate. In contrast, under the classic hindered diffusion model the flux was insensitive to the transition rate.

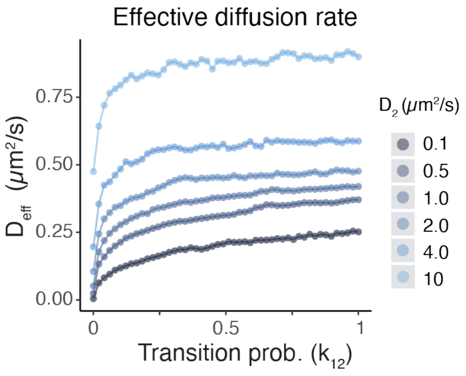


**Figure S11. Effective diffusion rate as a function of the diffusion rate of the unconstrained population ($D_2$)**

Under the topology-limited model, increasing the transition rate between constrained or free populations increased the effective secretion rate across a range of $D_2$ values. The constrained population was assumed to be constant at $D_1 = 0.5\mu m^2/s$ for all simulations.

**Movie S1.**

Representative movie of SHH-Halo diffusion, imaged with TIRF microscopy in the absence of SCUBE1. *(Top)* Raw images, scaled to 8-bit grayscale. *(Bottom*) Result of tracking with the TrackMate plugin in FIJI as described in supplemental methods. Images were acquired every 6ms, and played back in real time (166 Hz).

**Movie S2.**

SHH sender signaling to receivers without direct cell-cell contact in the presence of SCUBE1. mTurquoise2-CAAX; H2B-mTurquoise2-labeled SHH sender cells were plated with mCherry-CAAX; GBS-mCitrine receiver cells, in addition to unlabeled NIH3T3 cells, such that the imaging field is fully confluent but only sparsely plated with fluorescent cells, to enable cytoneme identification. 5% SCUBE1 conditioned media was added to the coculture. Images were acquired every 15 minutes for 62 hours.
